# Mapping RNA dependent RNA polymerase activity and immune gene expression using PRO-seq

**DOI:** 10.1101/2020.06.15.151738

**Authors:** Lakshmi S. Mahajan, Grace Eunyoo Kim, Hojoong Kwak

## Abstract

Positive strand, single strand RNA viruses ((+)ssRNA viruses) are viruses with an RNA genome that have broad impacts on a wide range of hosts, including SARS-CoV-2 human respiratory infections. Their replication and gene expression are driven by RNA dependent RNA polymerases (RdRp). Detecting active RNA synthesis by RdRp is critical for assessing the infectivity and pathogenicity of (+)ssRNA viruses. Current approaches rely on viral RNA detection, which cannot distinguish viral titer from RdRp activity. Precision Run-On sequencing (PRO-seq) is a nuclear run-on based nascent RNA sequencing method, widely used to map eukaryotic RNA polymerases by using labeled nucleotide analogues. Here we provide evidence that PRO-seq also detects RdRp activity and can serve as a highly sensitive RdRp mapping method. Coupled to PRO-seq in human blood samples, we propose to use PRO-seq as a single package method to detect (+)ssRNA virus RdRp activity and its interaction with host immune response through transcriptome-wide profiling of leukocyte gene expressions at once.

## Brief Communication

Positive strand, single strand RNA viruses ((+)ssRNA viruses) are viruses with an RNA genome that have broad impacts on a wide range of hosts, including coronaviruses in human respiratory infections^1,2^. Their replication and gene expression are driven by RNA dependent RNA polymerases (RdRp)^3,4^. Detecting active RNA synthesis by RdRp is critical for assessing the infectivity and pathogenicity of (+)ssRNA viruses. Current approaches rely on viral RNA detection, which cannot distinguish viral titer from RdRp activity^5^. For example, immune cell tropism of coronavirus is a factor related to the severity of the disease outcome currently under debate^6,7^, and it will be critical to distinguish the simple presence of viruses in blood from active viral expression inside blood cells.

PRO-seq is a nuclear run-on based nascent RNA sequencing method, widely used to map eukaryotic RNA polymerases (RNAP) by using labeled nucleotide analogs^8,9^. Here we provide evidence that PRO-seq can also detect RdRp activity, and propose that PRO-seq can serve as a highly sensitive RdRp mapping method. Coupled to PRO-seq in human samples, we anticipate the detection of RNA viruses and their activities through RdRp mapping. PRO-seq in blood samples of (+)-strand ssRNA virus infected patients will further define if there is immune cell tropism of the virus, and host-virus interactions: if the virus interferes with expression levels of the host factors, and if host immune gene expression affects the severity of the infection.

Drosophila A virus (DAV) is a (+)-strand ssRNA virus with low pathogenicity, infecting up to 30% of natural Drosophila populations^10^. We previously found that it is also transmitted in experimental Drosophila cell lines such as Snyder-2 (S2) cells. DAV is among the picornavirus family that encodes RdRp as the replicating and transcribing RNA polymerase^11^. Drosophila S2 cells were among the first cells to which PRO-seq methods were implemented to map RNAP in high resolution^8^. We reanalyzed the PRO-seq data in S2 cells, and investigated whether any of the PRO-seq reads were derived from non-Drosophila sequences. In the PRO-seq data with all 4 biotin-NTPs supplied in the Run-On reaction, 9.21 million reads were sequenced, 7.22 million reads (78%) were aligned at least once to the Drosophila melanogaster genome (Dm6), and 1.99 million reads were not mapped to the Dm6 genome. We re-aligned these unmapped reads to the collection of NCBI all virus sequences (12,442 virus refseq sequences), and 39,624 reads mapped to at least once virus sequence (2.0% of unmapped reads). 31,417 of those reads mapped to the DAV genome (NC_012958; 79% of viral sequences), indicating that PRO-seq reads from viral origin in S2 cells are predominantly from DAV. The PRO-seq reads from DAV covered the entire span of the DAV genome mostly on the plus strand (**Figure 1A**).

**Figure 1.**
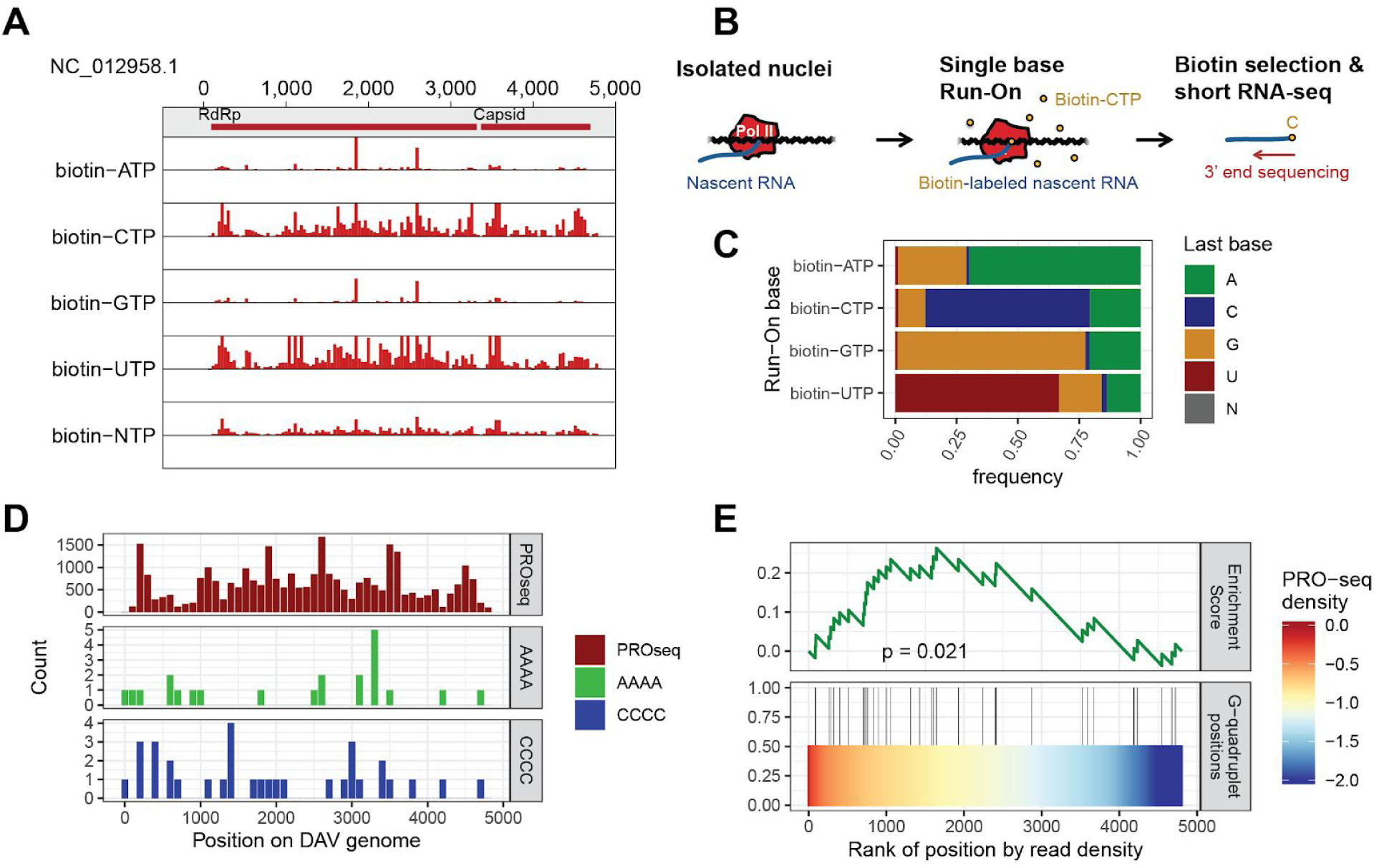
PRO-seq detects RdRp activity. **A.** PRO-seq read density profiles on the DAV genome. Both the plus (red) and minus (blue) strand reads are normalized to the same sequencing depths in all PRO-seq 5 libraries. Biotin labeled NTPs are listed along the left side of the figure, and position in bases is shown across the top of the figure. Red bars on the shaded top panel reflect ORFs for the RdRp and the Capsid proteins. Note that the PRO-seq reads from DAV cover the DAV genome on the positive strand almost entirely. **B.** Schematics of single-NTP PRO-seq (example of CTP). This image describes the process of labeling and sequencing biotin-labeled RNA transcripts, using CTP as an example. This process was repeated with ATP, GTP, CTP, and UTP. The first graphic (left) shows how in isolated nuclei, nascent RNAs are transcribed by the polymerase. The second graphic (middle) shows how biotin-CTP is introduced and incorporated into the nascent RNA to create biotin-labeled nascent RNA at which point transcription ends. The third graphic (right) shows how biotin-labeled RNA transcripts are sequenced from the 3’ end. **C.** Base frequency of DAV mapped reads in single-NTP PRO-seq libraries. The last base frequency is indicated along the left side of the figure, and the specific biotin labeled NTP is indicated along the bottom of the figure. The green portions indicate A as the last base, blue portions indicate C as the last base, yellow portions indicate G as the last base, red portions indicate U as the last base, and gray portions indicate N as the last base. In each case, approximately 65-70% of the PRO-seq reads that were mapped to the DAV genome ended with the same base as the run-on base. This indicates that most of the PRO-seq reads originating from the DAV genome are a result of RdRp activity. However, absolute mapped read numbers are higher in biotin-CTP and biotin-UTP libraries. **D.** PRO-seq density on DAV genome and base-quadruplet counts.The read count is indicated along the left side of each graph. The read densities for the PRO-seq data are much higher (some in the thousands) than the quadruplet read densities (less than ten). The first graph contains PRO-seq density data in red. The second graph shows A-quadruplet density data in green. The third graph shows C-quadruplet density data in blue. The position in bases is indicated along the bottom of each graph. Regions that show increased densities of PRO-seq are indicative of RdRp pausing. This can be seen at positions 200-300, 3,500-3,700, and 4,500-4,600. **E.** Enrichment analysis of template strand G-quadruplets on high PRO-seq density regions. G-quadruplets on the template strand are associated with RdRp stalling because C-quadruplets are more frequent in ORF-proximal PRO-seq peaks at the 200-300 base regions. The left side of each graph indicates read density. The bottom of the figure indicates the rank of each position by read density. In the bottom graph, G-quadruplet positions are mapped. Density score is 0.0 in the red region, -1.0 in the yellow region, and -2.0 in the blue region, as indicated by the key on the right-hand side. The enrichment score is graphed in the top graph in green, with a p-value of 0.021 indicating a significant difference in densities.

While PRO-seq is highly specific for nascent, biotin-labeled RNAs, it is also possible that DAV PRO-seq reads may have derived from the mature DAV RNA genome that contaminated the sample in small frequencies. The referenced PRO-seq study is originally designed to rule out this possibility by including Run-On libraries with one type of biotin-NTP supplied at once without the presence of any other NTP substrates. For example, PRO-seq biotin-CTP library is produced after washing out other NTP pools and supplying only the biotin-CTP in the Run-On reaction. This allows only the incorporation of biotin-CTP at the active site of the RNA polymerases if the RNA polymerase is positioned at the C base, because no other NTP substrates are available for the Run-On reaction. Therefore, 3′ ends of the PRO-seq RNA reads should predominantly be the C base (**Figure 1B**). In the original analysis, up to 90% of all reads ended with the supplied biotin-NTP base (A), supporting that PRO-seq detects nascent RNA transcribed by RNA polymerase. We performed the last base composition analysis on the PRO-seq reads aligned to the DAV genome (**Figure 1C**). Overall, 65-70% of the PRO-seq reads mapped to the DAV genome ended with the same base as the run-on base. If the reads were derived from RNA molecules in viral particles, the base composition would have been the same as the DAV genome composition (A:C:G:U = 29:24:24:24), which is not observed. This indicates that RdRp can incorporate biotin-NTPs similarly to nuclear run-ons, and the majority of PRO-seq reads from the DAV genome are the products of RdRp activity.

While all 4 single base run-ons show expected last base frequencies, the absolute amount of reads mapped to the DAV genome differs between each single base run-on (**Table 1, Figure 1A**). Biotin-CTP and biotin-UTP PRO-seq libraries show up to 10 times more reads mapped to the DAV genome than biotin-ATP and biotin-GTP libraries. This suggests that RdRp may be less efficient in using biotin-purine NTPs, and that biotin-pyrimidine NTPs (biotin-CTP and biotin-UTP) are the preferred substrates. This is different from Drosophila RNA Polymerase II, which appears to have comparable substrate specificity for all 4 biotin-NTPs.

**Table 1.** S2 PRO-seq statistics (re-analysis of data from Kwak *et al.*, Science 2013).

| Library | Reads | Aligned to Dm6 | Unmapped | Aligned to DAV |
| --- | --- | --- | --- | --- |
| PRO-seq 4biotin-NTP | 9,214,511 | 7,221,926 | 1,992,585 | 31,417 |
| PRO-seq biotin-ATP | 12,890,512 | 9,792,948 | 3,097,564 | 16,836 |
| PRO-seq biotin-CTP | 13,692,046 | 10,853,606 | 2,838,440 | 110,416 |
| PRO-seq biotin-GTP | 12,229,104 | 9,426,474 | 2,802,630 | 11,871 |
| PRO-seq biotin-UTP | 13,364,374 | 9,694,997 | 3,669,377 | 136,463 |

The PRO-seq reads cover the entire DAV genome, but some regions show increased PRO-seq densities suggestive of RdRp stalling or pausing. For example, there are increased read densities 100-200 bases downstream of the open reading frame (ORF) of RdRp (positions 200-300) and capsid protein of DAV (positions 3,500-3,700), reminiscent of promoter proximal pausing in eukaryotes. There are also read accumulations near the end of the ORFs (positions 4,500-4,600).

The ORF-proximal accumulation coincides with quadruplet rich regions of the template-strand of the DAV genome (**Figure 1D**). For example, C-quadruplets are more frequent in ORF-proximal PRO-seq peaks at 200-300 base regions (**Figure 1E**). This suggests the complementary G-quadruplets on the template strand are associated with RdRp stalling. G-quadruplets are known to form stable secondary RNA structures, and may provide physical barriers to RdRp elongation. Therefore, sequence features of the DAV genome can play a critical role in RdRp processivity, and these RNA sequence features can become therapeutic targets for clinical ssRNA virus infections.

We then examined if PRO-seq can be used to detect RdRp activity in human samples. We previously developed a PRO-seq procedure for a small amount of peripheral whole blood, which will detect the transcriptional landscape of nucleated leukocytes. This procedure uses biotin-CTP in the presence of 4 other NTPs as the substrates, and should be compatible with RdRp. One of the hypotheses on ssRNA virus infections is that they may drive immune cell response to aggravate symptoms of infection, possibly by directly infecting the immune cells.

For example, a recent study suggested that SARS-CoV-2 can directly infect T cells and induce T cell exhaustion^6^. PRO-seq in peripheral blood cells can not only test such hypotheses on viral activity by detecting RdRp, but also can provide comprehensive profiles of gene expression and transcriptional enhancer activities at the same time. This will allow us to examine host-ssRNA virus interactions with quantitative measures.

We analyzed 13 peripheral blood Chromatin Run-On (pChRO-seq) datasets, a modified version of PRO-seq, using 0.5 - 1.0 ml blood samples from 10 de-identified individuals, whose conditions regarding any viral infection were unknown. We collected PRO-seq reads that did not map to the Hg38 genome, and aligned them to the NCBI all virus database as described above. From this data, we did not detect significant PRO-seq sequences from (+)ssRNA viral genomes, indicating that none of the individuals had direct viral infections in the blood immune cells. For example, unlike the S2 cell data, we did not find any sequences specific to Drosophila A virus genome. This can serve as a baseline for evaluating human samples with active (+)ssRNA virus infections.

Viral infections lead to a diverse range of severity in clinical symptoms, and gene expression in immune cells such as cytokine response is an important factor in host-virus interaction. Our PRO-seq data show expression levels of immune-response related genes from human peripheral blood leukocytes. Out of 3,299 “immune” related genes classified by Gene Ontology, we found 990 differentially expressed genes in at least 1 individual (false discovery rate < 0.05, DESeq). Typically, PRO-seq levels are two or more fold higher in one or more individuals in these differentially expressed genes (**Figure 2A**). These expression patterns form clusters of immune related genes and clusters of individuals (**Figure 2B)**. The co-clustering of biologically replicated data from one individual (P1, P1n, P1nh) shows the reproducibility of PRO-seq in peripheral leukocytes (pChRO). The gene expression patterns suggest that individuals can be classified into groups with different immunity patterns if PRO-seq in peripheral leukocytes were conducted on a larger scale.

**Figure 2.**
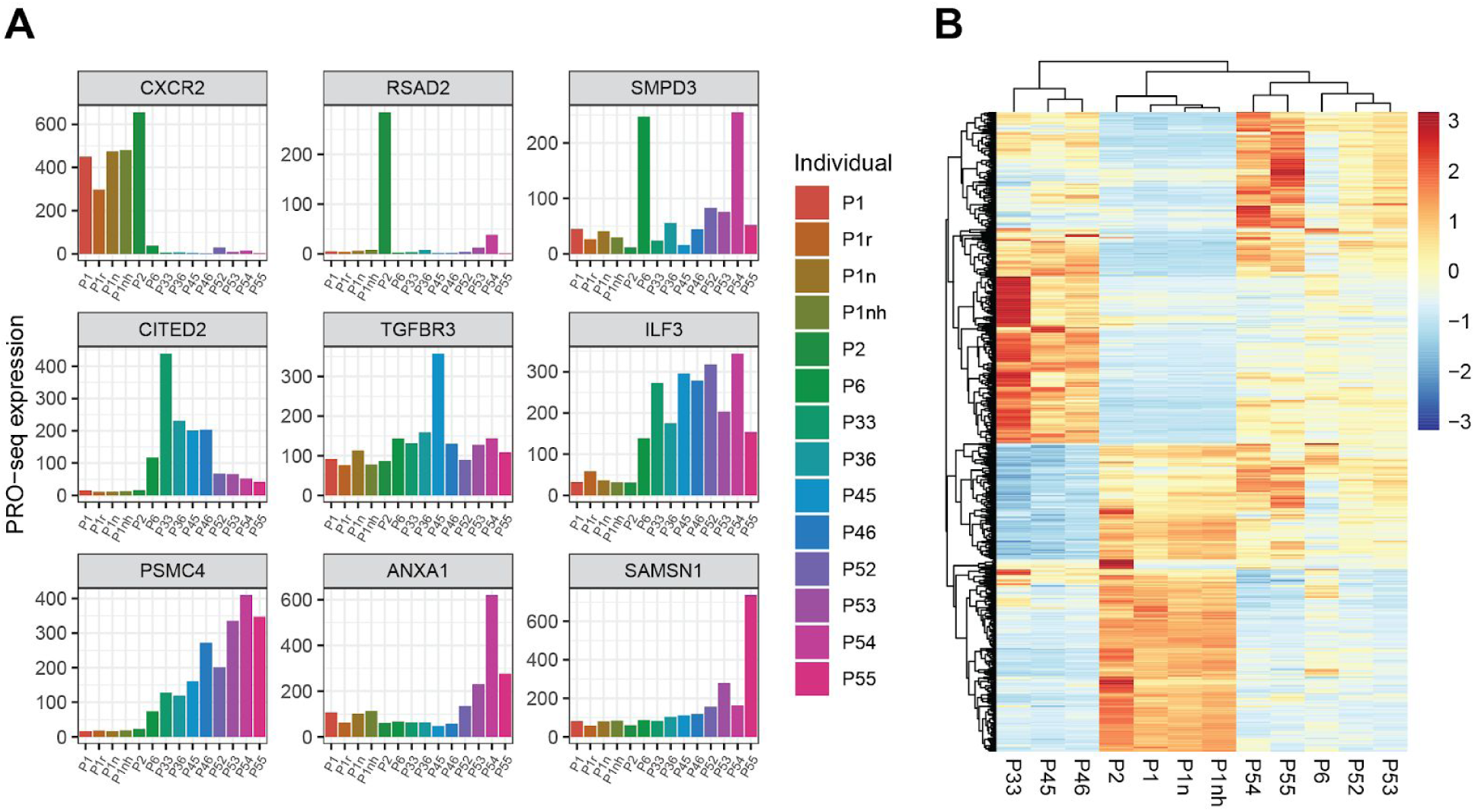
Transcriptome-wide profiles of leukocyte gene expression using PRO-seq. **A.** Expression levels of the most differentially expressed immune related genes in each individual. Expression levels were normalized using DESeq2, and the most differentially expressed genes were selected based on their p-values of the DESeq analysis. **B.** Heatmap of PRO-seq expression levels of the immune related genes and hierarchical clustering of the genes and individuals.Y-axis shows the 3,299 immune related genes clustered. Colormaps reflect the z-core normalized expression levels within the same genes across different individuals.

Overall, PRO-seq can serve as an extremely efficient method to both map RdRp activity and evaluate host immunity at once in human samples. The transcription/replication landscape of the (+)ssRNA viruses, as we report in DAV by PRO-seq, will provide a mechanistic basis of initiation and elongation by RdRp that can be directly applied to viruses that infect humans, such as SARS-CoV-2. RdRp is the direct target of the only approved drug for SARS-CoV-2 infection, Remdesivir, a chain terminating nucleoside analogue of RdRp. Identifying RdRp initiation and pausing sites in addition to defining sequence elements that regulate RdRp initiation and elongation can play a critical role in developing SARS-CoV-2 and other (+)ssRNA virus therapeutics.

Transcription/replication of the DAV genome by RdRp is not completely uniform throughout the whole genome, and there appear to be sites of RdRp slowdown, pausing, premature termination, and internal re-initiation. PRO-seq and 5′ end cap analysis of PRO-seq (PRO-cap) can precisely define these sites and sequence elements on the viral genome. Antisense RNA that interferes with these elements would be a straightforward therapeutic option that can synergize with nucleoside analog inhibitors such as Remdesivir. In addition, host factors that contribute to post-translational modification of eukaryotic transcription, replication, or RNA processing machinery can also play a role in RdRp processivity on the viral genome. Testing known small molecules using PRO-seq on a human cell based system, and directly measuring specific stages of RdRp initiation and elongation, will provide detailed information to identify drug candidates to be used in combination with other RdRp inhibitors. For example, secondary structures of the RNA genome may cause RdRp stalling as we observed in DAV, and drugs stabilizing RNA structures or inhibiting RNA helicases may also inhibit RdRp elongation. Such synergisms targeting RdRp from multiple mechanistic aspects will increase the efficiency of existing drugs such as Remdesivir and allow the use of lower tolerable doses in combinatorial therapies.

Finally, PRO-seq can provide measures of the expression levels of host factors while also measuring viral activity. While many (+)ssRNA viruses manifest as respiratory tract infections, there has been strong suspicion that severe cases may affect immune cells, either directly or indirectly. The versatility of PRO-seq to use blood leukocytes from untreated whole blood samples provides us with an opportunity to inspect the presence of actively infecting viruses in immune cells, and immune gene expression profiles that interact with viral infections at systematic levels at the same time. PRO-seq has the advantage of distinguishing inactive viral nucleic acids from actively transcribing and replicating viral genomes. Therefore, we propose to repurpose PRO-seq as an efficient strategy to map RdRp activities along with host gene expression profile to evaluate host-virus interactions simultaneously with high genomic resolution.

## Data accession

GEO SRX203291;SRR611824, SRR611825, SRR611826, SRR611827.

## Methods

### Identifying Drosophila A Virus (DAV) transcription from PRO-seq data

Drosophila PRO-seq sample sequences from a dataset included in Kwak et al, 2013 Science was retrieved from NCBI’s Gene Expression Omnibus (GEO) under the accession GSE42117. The samples are located in the Sequence Read Archive SRX203291. The four runs (SRR611824, SRR611825, SRR611826, SRR611827) were downloaded, adaptor sequences were removed using cutadapt, and the trimmed sequences were mapped to the reference Drosophila genome sequence (dm6) using STAR aligner. We aligned unmapped reads to the DAV sequence retrieved from the NCBI virus repository using the bwa bwa aligned.

### Human peripheral blood cell PRO-seq and analysis

Peripheral blood cell PRO-seq was performed as described previously^12^. Briefly, frozen blood sample is thawed and lysed in NUN buffer (0.3M NaCl, 1M Urea, 1% NP-40, 20mM HEPES, pH 7.5, 7.5mM MgCl2, 0.2mM EDTA, 1x protease inhibitor cocktail, 1 mM DTT, 4 u/ml RNase inhibitor) and chromatin is pelleted by centrifugation at 15,000 g for 20 min, 4°C. Pelleted chromatin was washed and used for the ultrashort PRO-seq procedure (uPRO). Chromatin or cells were incubated in the nuclear run-on reaction condition with biotin-NTPs and rNTPs supplied for 5 min at 37°C. Run-On RNA was extracted using TRIzol, and fragmented. 3′ RNA adaptors are ligated followed by 2 consecutive streptavidin bead bindings and extractions. Extracted RNA is converted to cDNA using template switch reverse transcription. After a SPRI bead clean-up, the cDNA is PCR amplified using primers compatible with Illumina Small RNA sequencing. Downstream analyses were performed as described previously^12^.

## References

1. Tay MZ, Poh CM, Rénia L, MacAry PA, Ng LFP. The trinity of COVID-19: immunity, inflammation and intervention. Nat Rev Immunol. 2020;20(6):363–374. doi: 10.1038/s41577-020-0311-8

2. Millet JK, Whittaker GR. Host cell proteases: Critical determinants of coronavirus tropism and pathogenesis. Virus Res. 2015;202:120–134. doi: 10.1016/j.virusres.2014.11.021

3. Gao Y, Yan L, Huang Y, et al. Structure of the RNA-dependent RNA polymerase from COVID-19 virus. Science. 2020;368(6492):779–782. doi: 10.1126/science.abb7498

4. Subissi L, Posthuma CC, Collet A, et al. One severe acute respiratory syndrome coronavirus protein complex integrates processive RNA polymerase and exonuclease activities. Proc Natl Acad Sci U S A. 2014;111(37):E3900–E3909. doi: 10.1073/pnas.1323705111

5. Chen W, Lan Y, Yuan X, et al. Detectable 2019-nCoV viral RNA in blood is a strong indicator for the further clinical severity. Emerg Microbes Infect. 2020;9(1):469-473. Published 2020 Feb 26. doi: 10.1080/22221751.2020.1732837

6. Wang X, Xu W, Hu G, et al. SARS-CoV-2 infects T lymphocytes through its spike protein-mediated membrane fusion [published online ahead of print, 2020 Apr 7]. Cell Mol Immunol. 2020;1–3. doi: 10.1038/s41423-020-0424-9

7. Chu H, Chan JF, Wang Y, et al. Comparative replication and immune activation profiles of SARS-CoV-2 and SARS-CoV in human lungs: an ex vivo study with implications for the pathogenesis of COVID-19 [published online ahead of print, 2020 Apr 9]. Clin Infect Dis. 2020;ciaa410. doi: 10.1093/cid/ciaa410

8. Kwak H, Fuda NJ, Core LJ, Lis JT. Precise maps of RNA polymerase reveal how promoters direct initiation and pausing. Science. 2013;339(6122):950–953. doi: 10.1126/science.1229386

9. Mahat DB, Kwak H, Booth GT, et al. Base-pair-resolution genome-wide mapping of active RNA polymerases using precision nuclear run-on (PRO-seq). Nat Protoc. 2016;11(8):1455–1476. doi: 10.1038/nprot.2016.086

10. Webster CL, Waldron FM, Robertson S, et al. The Discovery, Distribution, and Evolution of Viruses Associated with Drosophila melanogaster. PLoS Biol. 2015;13(7):e1002210. Published 2015 Jul 14. doi: 10.1371/journal.pbio.1002210

11. Ferrer-Orta C, Ferrero D, Verdaguer N. RNA-Dependent RNA Polymerases of Picornaviruses: From the Structure to Regulatory Mechanisms. Viruses. 2015;7(8):4438-4460. Published 2015 Aug 6. doi: 10.3390/v7082829

12. Kim SSY, Dziubek A, Lee SA, Kwak H. Nascent RNA sequencing of peripheral blood leukocytes reveal gene expression diversity bioRxiv 836841; doi: https://doi.org/10.1101/836841

